# Automated Quantitative Assessment of Elastic Fibers in Verhoeff–Van Gieson-Stained Mouse Aorta Histological Images

**DOI:** 10.1101/2025.05.01.651802

**Authors:** Austin E. Y. T. Lefebvre, Martin Mullis

## Abstract

The mechanical resilience of the aortic wall hinges on the organisation of its concentric elastic laminae, yet histological evaluation of these fibers remains largely qualitative and observer-dependent. We present a fully automated, stain-aware pipeline that transforms Verhoeff–Van Gieson (VVG) whole-slide images of mouse aortae into reproducible, quantitative maps of elastic-fiber architecture. Leveraging optical-density deconvolution to disentangle elastin from collagen, the workflow couples multi-resolution processing with graph-based skeletonisation to preserve gigapixel detail while scaling efficiently. It returns pixel-level measurements of fiber thickness, tortuosity, lamina count and network complexity, together with validation snapshots for transparent quality control.

By eliminating observer bias and delivering high-throughput morphometry, our framework enables powered genotype–phenotype screens in genetically diverse mouse populations and provides objective read-outs for interventions aimed at preserving matrix integrity. The modular codebase is open-source, readily extendable to other elastin-rich tissues or stains, and forms a bridge between qualitative microscopy and biomechanical phenotyping—setting the stage for large-scale, data-driven exploration of vascular structure–function relationships.

## Introduction

The mammalian aorta must repeatedly expand during systole and recoil during diastole, a feat made possible by the highly organized elastic laminae that dominate the tunica media^1^. In mice as in humans, these lamellae—concentric sheets of elastin interwoven with smooth-muscle cells—govern arterial compliance and dampen the pulsatile energy ejected by the heart^2,3^. Disruption of their architecture is a pathological hallmark of aneurysm, atherosclerosis, hypertension, and other vasculopathies^4–17^. In these conditions, the degradation or disorganization of elastic fibers can compromise the mechanical integrity of the aortic wall, leading to life-threatening consequences.

Despite their clinical importance, elastic fibers in histological sections are still assessed largely by eye. Traditional Verhoeff-Van Gieson (VVG) staining gives excellent contrast between elastin and surrounding tissue^18–22^, but the ensuing evaluations are qualitative, observer-dependent, and difficult to reproduce. Quantitative descriptors—fiber thickness, layer count, continuity, and waviness (tortuosity)—would offer far greater power to link microstructure with mechanical function or disease progression, yet remain under-utilized in routine practice.

Automated quantification is challenging for three reasons. First, VVG sections exhibit overlapping chromogens and uneven staining that obscure elastin boundaries. Second, the laminae form a branching, undulating network that resists simple morphological filters. Third, whole-slide images are large, demanding efficient algorithms that preserve fine detail while scaling to gigapixel data. Prior approaches address subsets of these problems but seldom capture the full spectrum of elastin morphology or handle high-throughput workflows.

Recent advances in computational pathology—optical-density (OD) deconvolution, graph-based skeletonization, and fast network analytics—make a comprehensive solution feasible. By coupling stain-aware segmentation with pixel-wise morphometry, a pipeline can now deliver reproducible, high-resolution maps of elastin thickness, continuity, layer number, and tortuosity across entire slides. Such data are invaluable for genotype–phenotype screens, longitudinal studies of disease models, and pre-clinical trials of matrix-targeted therapies.

Here, we present an end-to-end image-analysis framework that automatically extracts and quantifies elastic-fiber architecture from VVG-stained mouse aorta sections. The workflow (i) isolates elastin via OD deconvolution, (ii) converts complex laminae into a graph representation that retains branching topology, and (iii) reports a battery of morphological metrics together with intuitive validation images. By eliminating observer bias and standardizing measurements, our approach aims to accelerate vascular-biology research and provide a scaffold for future diagnostic or therapeutic applications.

## Results

### Image Preprocessing

Whole-slide images were acquired using high-resolution digital scanners as described in the methods section. To efficiently process large histological images and focus our analysis on relevant tissue regions, we implemented an automated ROI extraction procedure. The algorithm is implemented in Python and leverages several image processing libraries, including scipy, tifffile, skimage.morphology, and histo_tools.deconvolution^23–25^. The workflow consists of multiple steps, including downsampling, optical density (OD) calculation, background masking, ROI segmentation, and bounding box extraction, as outlined in Figure 1.

**Figure 1.**
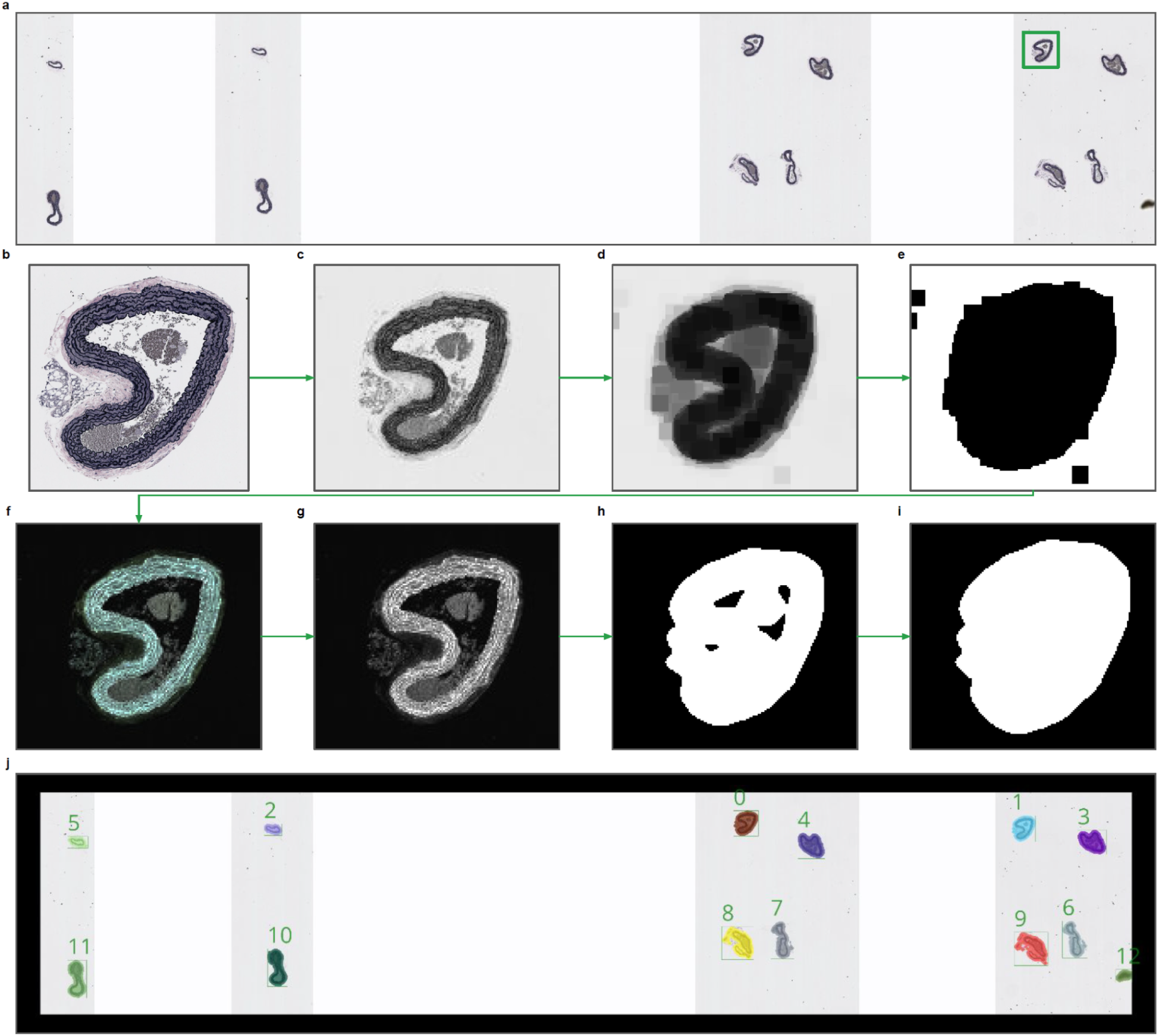
Automated Region of Interest (ROI) Extraction and Optical Density (OD) Processing Workflow. **a.** Whole-slide images (WSIs) were downsampled by a factor of 16 to balance computational efficiency and detail retention. **b.** The original downsampled RGB image was used for further analysis. **c.** The median intensity of the image across the RGB channels was computed for background masking. **d.** A minimum filter was applied to smooth the background and reduce noise in the median image. **e.** A background mask was generated to isolate non-tissue regions based on intensity thresholds. **f.** Optical density (OD) values were computed for each RGB channel using Beer-Lambert’s law. **g.** The maximum intensity projection of the OD image was calculated to highlight stained tissue regions. **h.** The initial binary mask of tissue regions was generated through Gaussian filtering and thresholding. **i.** Morphological operations refined the mask, filling holes and removing small irrelevant objects. **j.** Labeled ROIs were extracted, and bounding boxes were assigned to each detected region in both downsampled and full-resolution images.

Whole-slide images (WSIs) were first loaded using the tifffile library (Fig. 1a). The method get_downsample_multiplier() enables the selection of an appropriate downsampling factor by determining the closest precomputed resolution level in the multi-resolution pyramid of the WSI. Whole-slide images are typically stored in a pyramid format, where each successive level in the pyramid represents an image downsampled by a factor of 2 compared to the previous level. This allows for efficient viewing and processing of the image at various levels of detail. In this study, a downsample multiplier of 16 was chosen, meaning that the slide image was reduced by a factor of 16 along both the width and height dimensions. This results in an image that is significantly smaller and faster to process, but still sufficiently detailed to identify regions of interest (ROIs). Specifically, the function checks the size of the highest resolution (level 0, Fig. 1b) and compares it with the dimensions of the lower levels to find a precomputed downsampled version that matches the desired multiplier. Specifically, for each level, the method checks whether the downsampling factor between level 0 and the current level matches the specified multiplier. The method iterates through the available levels in the image and compares the size of each level to the full-resolution image (level 0). If a level exists that matches the target downsample multiplier (16 in this case), that level is selected. This level provides a precomputed downsampled version of the WSI, which is computationally efficient to load and process. If no precomputed level matches the specified multiplier, the code uses scipy’s zoom function to create a custom downsampled version. This is done by applying a scaling factor calculated based on the closest available resolution level. The scaling factor is computed as the ratio of the original image size to the desired downsample multiplier.

This zoom operation efficiently resizes the image to the required downsample level while maintaining the RGB structure of the image. The resultant image is stored in the variable im_rgb_downsample. Once the appropriate downsampled image is selected, the actual downsample scaling factor is calculated to ensure that any operations performed at this resolution can later be translated back to the original scale. This is critical when extracting regions of interest (ROIs) and calculating their coordinates in both the downsampled and full-resolution images. This scaling factor is stored in the downsample_scaling attribute and is used later to upscale the bounding boxes of detected ROIs from the downsampled image back to the full-resolution coordinates. A multiplier of 16 was chosen as it strikes a balance between computational efficiency and maintaining sufficient detail for region detection. At this downsampling level, the size of the image is reduced enough to allow for fast processing, but the resolution is still high enough to capture the key morphological features of the tissue, such as boundaries between different tissue types and the presence of structures of interest (e.g., glands, vessels). This is particularly important for algorithms that rely on feature extraction at a cellular or tissue level, as excessive downsampling could result in the loss of crucial information.

Optical density (OD) is a critical step in this workflow for improving the contrast between stained tissue and background, which facilitates the accurate detection of regions of interest (ROIs). OD calculation is based on the physics of light absorption in stained histological samples, where the relationship between the incident light, transmitted light, and the staining properties of the sample can be modeled using Beer-Lambert’s Law. In histological images, tissue sections are often stained with dyes (e.g., hematoxylin and eosin) to highlight different structures. The color intensity in RGB images correlates with the concentration of these stains. However, direct use of raw RGB values can be unreliable for segmentation due to variability in illumination and staining intensity. By converting RGB values into OD values, we can normalize these variations, making it easier to differentiate tissue regions from the background and isolate features of interest. The OD calculation follows a multi-step process, beginning with background masking to correct for background illumination, followed by conversion of the raw RGB values into OD values using a logarithmic transformation. Specifically, the background mask is essential to isolate non-tissue regions, which typically correspond to white or near-white areas in the image (i.e., glass slide or unstained regions). This step helps correct for any residual background illumination or staining artifacts. The background mask is calculated by first taking the median pixel intensity across the RGB channels of the downsampled image (Fig. 1c). Then, a minimum filter (with a size of 10 pixels) is applied to smooth the background and remove noise (Fig. 1d). Finally, the median intensity of the image is used to identify pixels with values in the range of 220 to 245, which corresponds to near-white regions typically considered as background, and is used as a threshold (Fig. 1e). This results in a binary mask of the background, which is used to exclude these areas in the subsequent OD calculations. Once the background mask is obtained, the next step is to estimate the incident light intensity (I0), which represents the theoretical maximum light intensity in unstained regions. This is necessary for calculating the OD image using Beer-Lambert’s Law. The function deconvolution.get_i0() is used to compute I0, by averaging the pixel values of the background regions (masked out by the background mask) across the RGB channels. This provides a set of I0 values, one for each of the RGB channels, which will serve as the reference for the OD calculation. With the incident light intensity I0 estimated, we can now calculate the OD values for each pixel in the image using the Beer-Lambert Law:

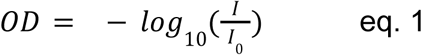

Where *I* is the pixel intensity in the RGB channels of the tissue image, *I*_0_ is the incident light intensity for each channel, calculated from the background regions. The function deconvolution.get_od() applies this logarithmic transformation to convert the RGB values into OD values. This transformation normalizes the effect of illumination variability and accentuates differences in stain absorption, which is important for accurately identifying tissue regions. Since logarithmic operations can lead to infinite or NaN values (especially when *I* approaches 0), any resulting infinite or NaN values in the OD image are replaced with zeros. The resulting OD image is a three-channel (RGB) image, where each channel represents the optical density for the corresponding RGB channel (Fig. 1f). For region detection, the maximum projection of the OD image is calculated along the RGB channels to create a single grayscale OD image, which emphasizes the most prominent stain absorption across the channels (Fig. 1g). This OD image is now ready for subsequent thresholding and region detection, as it enhances the contrast between stained tissue and background areas, making it easier to isolate regions of interest (ROIs).

The purpose of ROI detection and masking is to identify regions of interest (ROIs) within the optical density (OD) image that represent significant tissue structures. The masking step leverages the enhanced contrast from the OD calculation to segment meaningful tissue areas while filtering out background and irrelevant regions. Once the OD image is calculated, the next step is to threshold the image to isolate potential regions of interest (ROIs) based on intensity. The OD image, being a grayscale projection of the original RGB image, highlights stained tissue regions with higher intensity values, while the background and unstained areas are typically darker. A Gaussian filter is applied to smooth the OD image and reduce noise before thresholding. This is necessary because raw OD images can contain small variations or artifacts that could result in false-positive detections if thresholded directly. A filter with a sigma value of 3 is applied to blur the image, which helps smooth out small, insignificant variations and makes the subsequent thresholding more robust. A threshold value (od_threshold) of 0.1 is applied to the smoothed OD image. This threshold effectively distinguishes between tissue regions (above the threshold) and background (below the threshold). The result is a binary mask where pixels corresponding to tissue regions are marked as True (1), and background pixels are marked as False (0). This forms the initial segmentation mask (see Fig. 1h). After thresholding, the resulting binary mask may contain small holes or gaps within the detected tissue regions due to noise, imperfections in staining, or the thresholding process itself. To ensure contiguous regions are detected, hole filling is performed using scipy.ndimage.binary_fill_holes(). This operation ensures that any internal holes in the binary mask, which are fully surrounded by tissue pixels, are filled in, creating a more continuous segmentation. This improves the integrity of the detected regions and ensures that they can be treated as cohesive objects. To further enhance the segmentation, a binary dilation operation is applied to the mask. This step is important because, in some cases, tissue regions may have thin boundaries or gaps that are not adequately captured during thresholding. Dilation expands the boundaries of the detected regions slightly, helping to bridge small gaps between neighboring regions. The dilation is repeated for a fixed number of iterations (5 in this case), ensuring that even small separations between nearby tissue regions are closed, improving the final segmentation mask’s robustness. Once the binary mask is dilated, small irrelevant objects (e.g., noise or tiny tissue fragments) need to be removed to clean up the segmentation. The function skimage.morphology.remove_small_objects() is used to filter out objects that are smaller than a specified size threshold, defined by the downsample scale. The default size is set to downsample_scaling/16 * 1000, ensuring that only regions larger than this threshold are retained. This prevents small specks or noise from being included as valid ROIs. This operation produces the final ROI mask, which accurately segments all the meaningful tissue regions while discarding small, irrelevant objects (Fig. 1i).

Once the ROI mask is created, the next step is to identify each distinct region (ROI) within the binary mask, assign labels to them, and extract their bounding boxes. This information is essential for indexing the regions and determining their spatial locations in both the downsampled and original image coordinates. The function scipy.ndimage.label() is used to identify and label connected components in the binary ROI mask. Connected components are groups of adjacent pixels marked as True (1) in the binary mask, which represent distinct tissue regions. This output is a labeled image, where each connected component (or tissue region) is assigned a unique integer label. Pixels in the binary mask corresponding to the first detected region are labeled as 1, the second region as 2, and so on. This labeled image is used to differentiate between the distinct regions of interest, each of which can be analyzed separately. Once the regions are labeled, the next step is to extract their bounding boxes. A bounding box is a rectangular region that fully encloses a detected ROI, defined by the coordinates of its top-left and bottom-right corners. The function scipy.ndimage.find_objects() is used to locate the bounding boxes for each labeled component. This function returns a list of slices for each labeled component. Each slice corresponds to a rectangular bounding box that contains the ROI. These slices are used to calculate the coordinates of the bounding box for each region. Bounding boxes are extracted both in the downsampled image and upscaled to the original resolution to ensure that ROIs can be mapped back to the full-resolution image. The bounding box coordinates are calculated directly from the slices returned by find_objects(). These coordinates represent the region in the downsampled image. To map the bounding boxes back to the original full-resolution image, the coordinates are multiplied by the downsample scaling factor (self.downsample_scaling). This ensures that the bounding boxes are accurately translated to the original resolution. The bounding box coordinates in both resolutions are stored as lists, allowing for downstream applications such as displaying the ROIs or analyzing the regions in their full-resolution context. Finally, the labeled image (label_im), the list of bounding boxes in the downsampled image (region_bounding_boxes), and the upscaled bounding boxes (region_bounding_boxes_upscaled) are stored as attributes of the Slide object. These stored attributes are available for subsequent analysis, visualization, or region-specific operations (Fig. 1j).

### Elastin Segmentation and Network Construction

The optical density (OD) image, im_od, is critical in the detection of elastin layers and their structures within the tissue. First, the region of interest (ROI) is extracted from the whole-slide image (WSI) based on the bounding boxes, which represent areas of interest. This is done in the get_full_rgb_im() method, where the RGB image for the bounding box is retrieved from the full-resolution slide (Fig. 2a). The resulting RGB image is then used as input for OD processing. To generate the OD image, the deconvolution method is applied to the extracted RGB image. This process separates the VVG stain channels from the RGB image, improving the detection of specific elastin structures. The OD deconvolution removes any artifacts from poorly stained regions by applying a threshold to mask pixels where the intensity is too high (which usually corresponds to unstained or empty regions). This is done by calculating ratios between the RGB channels and applying a mask to remove unwanted background areas. First a maximum intensity projection across the RGB channels is produced (Fig. 2b). A mask is created to filter out regions where the maximum intensity across the RGB channels is too high, indicating that they are not part of the tissue (likely background or overexposed areas). The background mask is refined by Gaussian filtering the ratio between certain RGB channels (zero_two_mask and one_two_mask) to ensure only well-stained regions remain (Fig. 2c). The method deconvolution.run_full() is then applied, which uses a color deconvolution algorithm to separate the RGB channels into their respective stain components. This process results in the optical density (OD) image, im_od, which highlights the stained tissue structures, particularly the elastin layers (Fig. 2d). To isolate the regions containing tissue from the background, a tissue mask is generated from the OD image. This is done by analyzing the maximum projection of the OD image across the relevant channels (channel 0 and channel 1), which captures the strongest signal from the stained regions.

**Figure 2.**
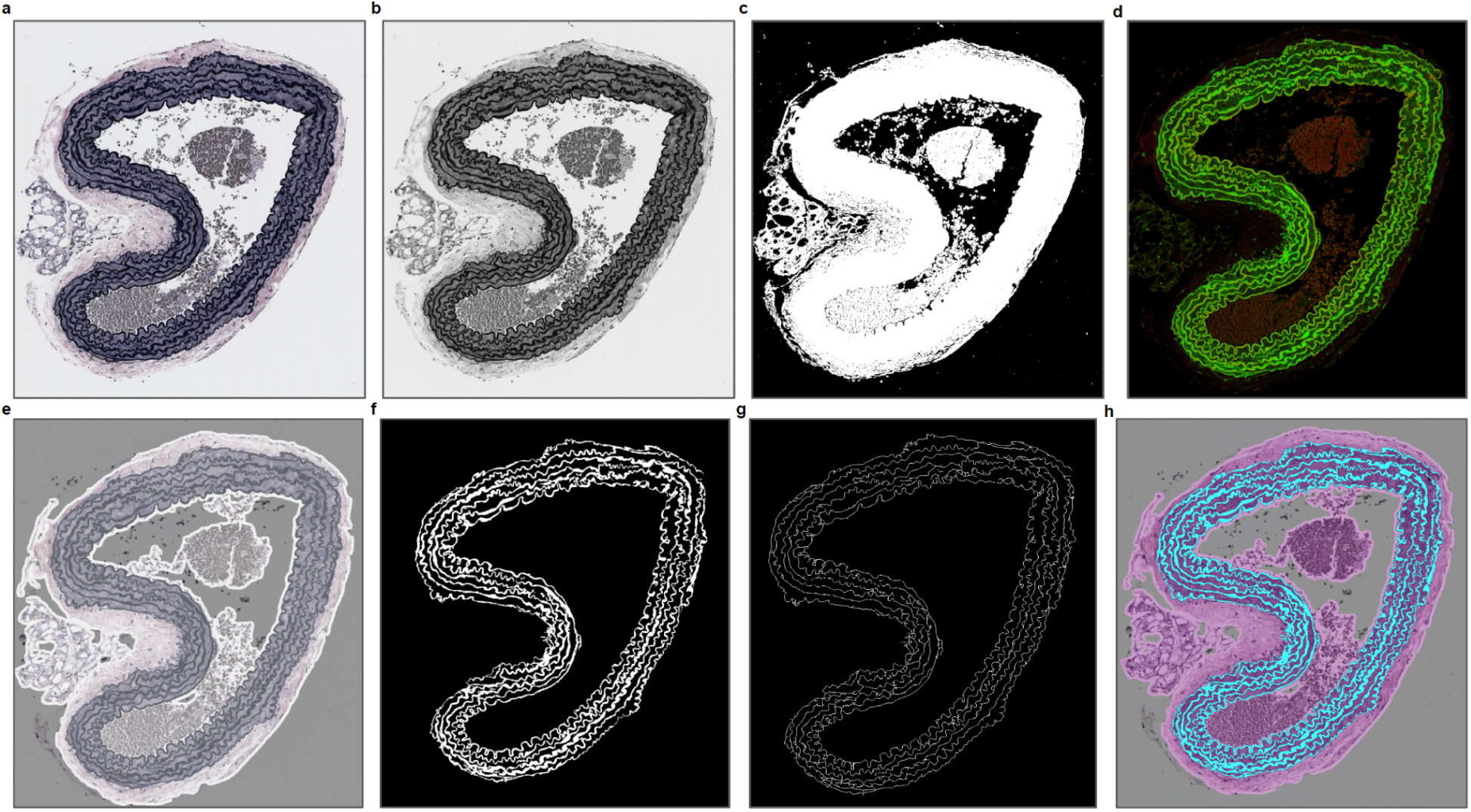
Elastin Segmentation and Network Construction Workflow. **a.** The full-resolution RGB image is extracted from the region of interest (ROI) based on the bounding boxes. **b.** A maximum intensity projection is created across the RGB channels, highlighting stained regions for further processing. **c.** A background mask is generated to remove overexposed or non-tissue areas, refining the tissue areas for elastin segmentation. **d.** The optical density (OD) image is generated using deconvolution, isolating stained elastin structures for further analysis. **e.** A binary tissue mask is created by thresholding the OD image, identifying regions of tissue while excluding background areas. **f.** The elastin mask is generated by thresholding the second channel of the OD image, isolating regions with high elastin density. **g.** Skeletonization is applied to the elastin mask, reducing the elastin regions to their centerlines, creating a simplified representation of the elastin network. **h.** The ratio of elastin to total tissue is calculated, visualizing the fraction of elastin present within the tissue slice.

The tissue mask is generated by thresholding this projection to retain only areas with sufficient intensity, which correspond to actual tissue. A Gaussian filter smooths the image and reduces noise. This filtered image then has small objects removed. Finally a uniform filter further smooths the mask to ensure that only large, contiguous areas of tissue are detected. The final result is a binary tissue mask (im_tissue_mask), where areas containing actual tissue are marked as True (1) and the background is False (0) (Fig. 2e).

Once the OD image is generated and the tissue regions are identified via the tissue mask, the next step is to specifically isolate the elastin layers and generate their skeleton. The elastin mask is obtained by thresholding the second channel of the OD image (which corresponds to elastin staining) to isolate regions where elastin is present. A binary mask is created by applying a threshold (0.8 in this case), which marks areas with high elastin density. This initial mask is then refined through several steps. A Gaussian filter is applied to the mask to smooth the boundaries and reduce noise. This is important to create a cleaner segmentation of elastin regions. To ensure that the elastin regions are contiguous and fill in small gaps, a binary closing operation is applied. This helps close small holes and connects nearby elastin regions. Small objects that do not represent meaningful elastin structures are removed using skimage.morphology.remove_small_objects(). This operation helps clean up the mask and ensures that only significant elastin regions are retained. The result is a binary elastin mask, im_elastin_mask, where regions containing elastin are marked as True (1) and the background is False (0) (Fig. 2f).

To further analyze the structure of the elastin fibers, the skeleton of the elastin mask is computed. Skeletonization reduces the elastin regions to their centerlines, which helps in measuring structural properties such as length, connectivity, and tortuosity. The skeletonization process is performed using the skimage.morphology.skeletonize() function, which creates a binary image where the elastin fibers are reduced to single-pixel-wide lines representing the centerline of the elastin structures (Fig 2g). This skeleton image (im_skeleton) is a simplified representation of the elastin fibers and is used for further analysis, such as calculating the tortuosity and other network properties. The ratio of the elastin mask pixels to the tissue mask pixels can also be used to calculate the fraction of elastin within the tissue slice (Fig. 2h).

### Elastin Thickness Extraction

The measurement of elastin thickness is a key component of analyzing the structural properties of elastin fibers within the tissue. Thickness refers to the distance between the outer boundaries of the elastin layers and is an important parameter for understanding the integrity and characteristics of the elastic lamina. In this section, we describe how the algorithm computes elastin thickness using the elastin mask and skeleton. The thickness of the elastin layers is computed using a distance transform on the elastin mask. The distance transform assigns each pixel in the elastin region a value corresponding to its Euclidean distance from the nearest background pixel (i.e., the boundary of the elastin region). This value represents the shortest distance from any point inside the elastin region to its edge. The distance transform is performed using scipy.ndimage.distance_transform_edt(), which calculates the Euclidean distance from each pixel in the elastin mask to the nearest zero (background) pixel. This generates a distance map (im_distance), where the pixel values represent the distance from the nearest boundary of the elastin region. In this distance map, pixels near the edge of the elastin region have smaller values and pixels near the center of the elastin region have larger values, representing the distance to the closest boundary (Fig 3a). The distance values from the distance map are used to compute the thickness of the elastin layers by sampling the distance values at the locations of the elastin skeleton pixels. The skeleton pixels are assumed to be located at the center of the elastin layers, and the distance from these pixels to the boundary of the elastin region gives an approximation of the half-thickness of the elastin layer (Fig. 3b). To obtain the full thickness, the distance values at the skeleton pixels are multiplied by 2. This is because the skeleton represents the midpoint of the elastin layer, and the distance transform provides the distance from this midpoint to the nearest boundary, so multiplying by 2 gives the total thickness from one boundary to the opposite boundary. The elastin thickness map is created by multiplying the skeleton by the corresponding distance transform values.

**Figure 3.**
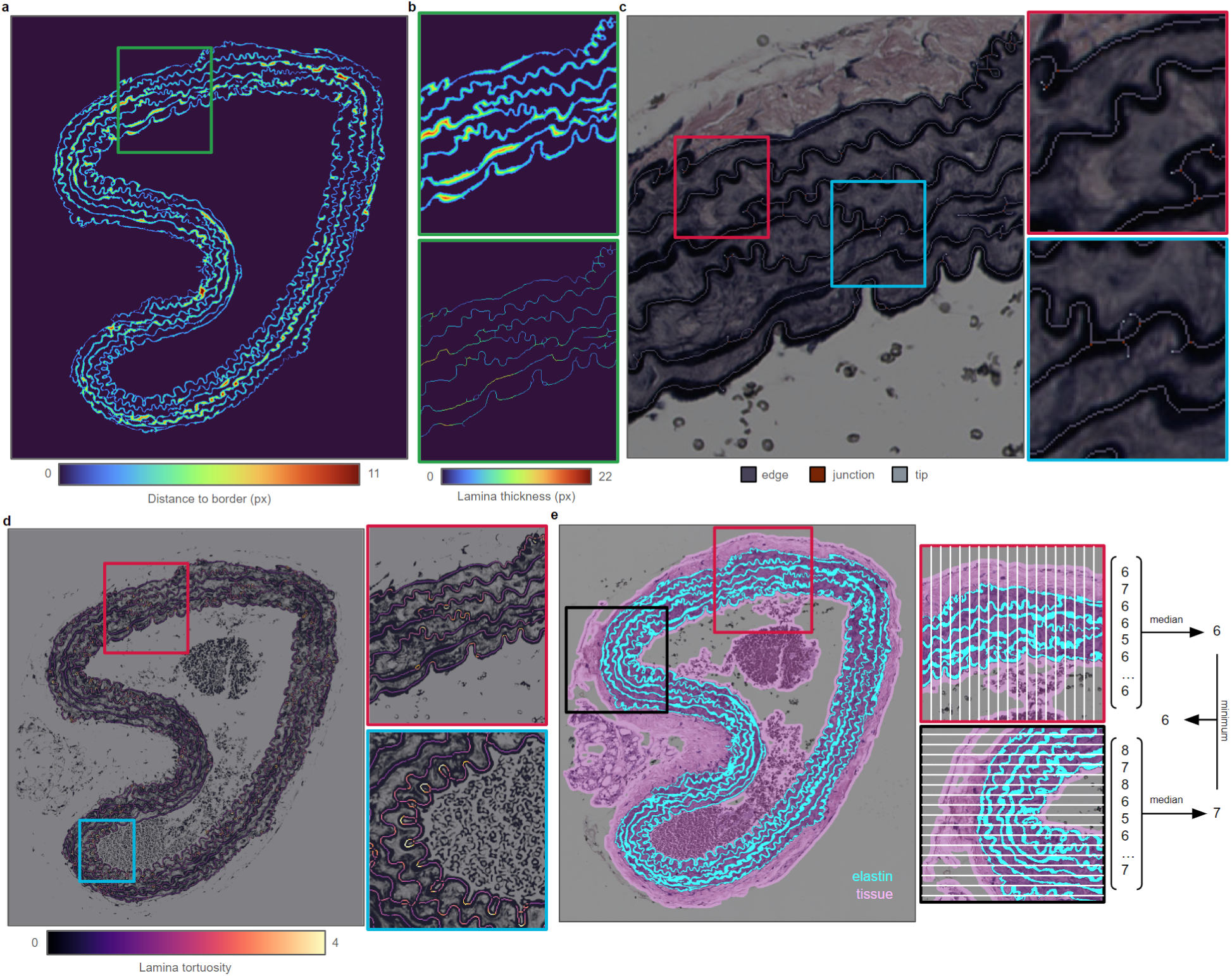
Elastin Thickness, Network, and Layer Number Extraction. **a.** Distance map showing the Euclidean distance from each pixel in the elastin region to the nearest boundary, used for calculating elastin thickness. **b.** The elastin thickness map generated by multiplying the distance map by the elastin skeleton, providing a pixel-wise representation of elastin thickness. **c.** The classification of network nodes into tip points (end of fibers), junctions (branch points), and edges (regular points) within the elastin network. **d.** Tortuosity map, visualizing the degree of curvature or waviness of the elastin fibers across the tissue, where higher values indicate greater curvature. **e.** Visualization of elastin layers within a tissue section, with row-wise and column-wise analyses to count distinct elastin layers, estimating the number of elastin layers in the tissue.

This results in an image, im_thickness_map, where each pixel in the skeleton has a value corresponding to the thickness of the elastin layer at that point. The elastin thickness map provides a detailed, pixel-wise representation of the thickness of elastin layers across the entire region. The thickness map can be visualized to identify regions where elastin layers are thin or thick, which may indicate areas of structural integrity or damage. The thickness map can be analyzed to compute statistical metrics, such as the mean, median, standard deviation, and range of elastin thickness values across the tissue.

### Elastin Graph Network

The elastin network represents the interconnected structure of elastin fibers, derived from the skeletonized elastin mask. This network is essential for analyzing the connectivity, complexity, and geometric properties of the elastin fibers, such as tortuosity and branching behavior. The network is built from the skeleton image (im_skeleton), which is a one-pixel-wide representation of the centerlines of the elastin fibers. In the skeleton, each pixel represents a point along the elastin fiber, and the network is constructed by identifying and connecting these points based on their spatial proximity. The Network class is responsible for constructing the elastin network. The network is initialized by passing the skeleton image to the constructor, where the pixel coordinates of the skeleton points are extracted. The coordinates of all non-zero pixels (skeleton points) in the skeleton image are extracted using np.where(). These coordinates represent the spatial positions of the points that make up the elastin network. Each skeleton pixel is represented as an individual node in the network, which is stored as a Pixel object. The Pixel class defines the properties of each node, such as its coordinates, the list of connected pixels, its node type (tip, branch, or regular), and the tortuosity of the fiber at that point. The skeleton coordinates are converted into Pixel objects using the get_pixels() method. This method iterates over the list of skeleton coordinates and creates a Pixel object for each coordinate. Each pixel in the skeleton is now represented as a node in the network, with properties such as coord (its location) and connected_pixels (a list of other pixels it is connected to). The next step is to define the connections between adjacent pixels in the skeleton. Pixels are connected if they are adjacent to each other in the image (i.e., within a 3x3 neighborhood). These connections form the edges of the network, linking adjacent pixels along the elastin fibers. There are two methods provided for establishing the connections between pixels:

### Method 1: Fast Connection Mask (get_connection_mask())

The method get_connection_mask() creates a connection matrix between pixels using a fast vectorized approach. This method constructs a location mask (loc_mask), which identifies pairs of pixels that are spatially close (within a distance of 1 pixel in the x and y directions) and should be connected. This mask is a boolean matrix where each entry indicates whether two pixels should be connected based on their proximity. The diagonal of the mask is set to False to prevent pixels from being connected to themselves. After constructing the connection mask, the method identifies pairs of connected pixels and updates the connected_pixels list for each Pixel object. For each pixel pair that is adjacent, the method appends the connected pixel to the connected_pixels list for both pixels.

### Method 2: Neighborhood Search (get_connection_mask_2())

An alternative, slower method, get_connection_mask_2(), is used if memory constraints prevent the fast connection mask from being applied. This method manually searches for neighbors within a 3x3 neighborhood around each pixel. For each pixel, the method checks all possible neighboring coordinates in a 3x3 grid around the pixel. If a neighboring pixel is found in the skeleton, it is added to the list of connected pixels.

Once the connections between pixels are established, the algorithm classifies each pixel into one of three node types: tip points, branch points, or regular points, based on the number of connected pixels each pixel has (Fig. 3c). The classification of node types is done in the set_type() method. If a pixel has exactly one connection, it is classified as a tip point, representing the end of a branch. If a pixel has three or more connections, it is classified as a branch point, where multiple fibers diverge or converge. If a pixel has exactly two connections, it is classified as an edge, or regular, point, representing a point along a fiber between two other points. In some cases, branch points may be connected to other branch points. These connections can be redundant or incorrect, so the method clean_branch_points() ensures that each branch point is appropriately connected by converting unnecessary branch points into regular points. This ensures that branch points are correctly identified and connected, and unnecessary branching is avoided.

After constructing the network and classifying the node types, several network statistics are calculated in the get_network_stats() method. The total number of branch points and tip points are counted. The number of distinct connected components (trees) in the skeleton is calculated using scipy.ndimage.label(). The total number of segments in the network is the sum of branch points and trees. The Segment-to-Network ratio is calculated by dividing the number of segments by the total length of the network. A lower value indicates more breaks in the network, while a higher value indicates greater continuity. The complexity of the network is a sum of the total segments and tip points, giving a measure of the overall intricacy of the network structure.

### Elastin Lamina Tortuosity

Tortuosity is a measure of the curvature or waviness of a path, and in this context, it refers to the degree of bending or undulation of the elastin fibers in the network. The tortuosity of each fiber is calculated at each pixel where the fiber is continuous (i.e., at regular points with two connections). The goal is to determine how much the fiber curves over a certain distance (defined by a search radius) and compare this curved distance to the straight-line distance between two endpoints. Tortuosity is only calculated for regular points (pixels where len(connected_pixels) == 2). These points are located in the middle of a segment of the fiber, where the fiber continues in two directions. The tortuosity calculation is performed by exploring the connections from each regular point in both directions to measure how the fiber curves. The algorithm checks whether a pixel has exactly two connected pixels, meaning it is a regular point and not a branch or tip. Tortuosity is calculated only for these points. To calculate tortuosity, the algorithm performs a depth-first search (DFS) starting from the regular point, exploring both directions of the connected fiber to locate the endpoints of the fiber segment within a specified distance. This distance is referred to as the search radius (search_rad), which limits how far along the fiber the search proceeds. The method undulation_depth_first() recursively explores connected pixels until the total distance traveled along the fiber reaches or exceeds the search radius. The search continues along connected pixels, adding up the Euclidean distance between each pixel and its connected neighbor. The DFS halts when the total distance traveled reaches or exceeds the specified search radius. At this point, the pixel is marked as an endpoint for this direction of the fiber. Two depth-first searches are performed, one for each direction along the fiber, starting from the current pixel. The result is two sets of endpoints (end_points) for the two directions, containing the pixels that are located at the ends of the fiber segment being examined.

Once the two endpoints are found, the next step is to calculate the tortuosity. Tortuosity is a measure of how much longer the fiber path is compared to a straight line between the two endpoints. To compute this, the algorithm first calculates the straight-line distance (Euclidean distance) between the two sets of endpoints. The x and y coordinates of the endpoints in both directions are collected into separate lists (dirs_x and dirs_y). A tortuosity matrix is then constructed by computing the Euclidean distance between all combinations of endpoints in the two directions. This matrix contains the Euclidean distances between each pair of endpoints found in the two opposite directions. The matrix allows the algorithm to determine the straight-line distance between the most distant points in both directions. Next, the curved distance is calculated by summing the pixel-to-pixel distances along the fiber. This was already done during the DFS, where the total distance traveled along the curved path was accumulated. Once the curved and straight-line distances are known, the tortuosity is calculated using the formula:

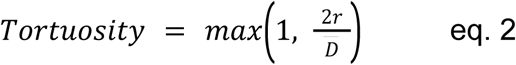

Where *r* is the search radius and *D̅* is the mean straight-line distance between all found end points. This formula compares the curved path (approximated by the search radius, which represents half the fiber segment being examined) to the mean straight-line distance between the endpoints. If the straight-line distance is much shorter than the curved path, the tortuosity will be greater than 1. If the straight-line distance is equal to or longer than the curved path, the tortuosity will be close to 1, indicating a relatively straight fiber. If the tortuosity matrix contains valid distances, the tortuosity value is calculated by dividing twice the search radius by the mean straight-line distance between the endpoints. The max(1.0, …) function ensures that tortuosity values cannot be less than 1.0 (since a perfectly straight line has a tortuosity of 1). If no valid endpoints are found (e.g., due to a short segment or incomplete fiber), the tortuosity is set to NaN (not a number). The tortuosity value for each regular point is stored in the tortuosity attribute of the corresponding Pixel object. Once the tortuosity values for all regular points are calculated, the method get_tortuosity_im() generates a tortuosity map (im_tortuosity), which visualizes the tortuosity of the elastin network across the entire region (Fig 3d). This map has the same dimensions as the skeleton image, and each pixel in the skeleton is assigned its corresponding tortuosity value. Pixels that are not part of the skeleton or do not have valid tortuosity values are set to zero. The tortuosity map is useful for visualizing how wavy or curved the elastin fibers are across the entire image. Areas with higher tortuosity values indicate regions where the fibers are more curved, while areas with values close to 1 indicate straighter fibers. The tortuosity map is also used for quantitative analysis, where various statistical metrics (mean, median, standard deviation, etc.) are computed to summarize the overall tortuosity of the elastin network. These metrics are stored in the tortuosity_stats attribute of the VVG class and can be used for further reporting or analysis.

### Elastin Layer Number

The number of elastin layers can also be an interesting metric to study, specifically estimating the number of distinct elastin layers in a given region of the tissue. The method works by analyzing the elastin mask derived from the optical density (OD) image and counting the number of distinct elastin components (or pieces) within each tissue section, both in horizontal (row-wise) and vertical (column-wise) directions. Before determining the number of elastin layers, the elastin mask (im_elastin_mask) and tissue mask (im_tissue_mask) must be generated. These masks serve as the foundation for identifying and counting elastin layers. The elastin mask is created by thresholding the second channel of the OD image (which represents elastin staining) to isolate areas containing elastin. Morphological operations are then applied to clean up the mask, as described in previous sections. This binary mask highlights the regions where elastin is present (Fig. 2f). The tissue mask is generated from the OD image to identify the areas containing actual tissue, excluding the background (Fig. 2e). This ensures that only the elastin layers within the tissue are counted. The number of elastin layers will be determined by examining the elastin mask within the tissue regions defined by the tissue mask. The first step in determining the number of elastin layers is to label the distinct tissue regions within the tissue mask. This is done using scipy.ndimage.label(), which assigns a unique label to each connected component (or object) in the binary tissue mask. This is a labeled image where each distinct piece of tissue is assigned a unique integer label. Each labeled region corresponds to a connected area of tissue where elastin layers may be present.

For each labeled tissue region, the algorithm analyzes the intensity of the elastin mask to count how many separate elastin components are present. This is done by analyzing the elastin components both row-wise (horizontally) and column-wise (vertically) within the region (Fig. 3e). The scipy.ndimage.label() function is applied again, this time to the elastin mask within each row and column of the tissue region. This function counts the number of distinct connected elastin pieces (or chunks) in each row or column, which corresponds to the number of elastin layers intersecting that row or column. Here, regionprops extracts the properties of the labeled tissue regions, including their corresponding portions in the elastin mask (obj.image_intensity). The method then proceeds to count the number of elastin layers in each row and column for each tissue object. For each tissue object, the algorithm iterates through every row and column of the elastin mask within the object, counting the number of distinct elastin pieces (or layers) that appear in that row or column. The algorithm starts by analyzing each row in the elastin mask of the tissue object. For each row, it uses scipy.ndimage.label() to find connected components (elastin chunks) in that row and counts the number of elastin layers in each chunk. After completing the row-wise analysis, the same process is applied column-wise to count the elastin layers in each column of the elastin mask for the tissue object. After counting the number of elastin layers in both rows and columns, the final number of elastin layers for the tissue object is determined by taking the minimum of the median number of elastin layers from the row-wise and column-wise lists. The median number of elastin layers from the row-wise and column-wise analyses gives a robust estimate of the typical number of layers within the tissue object. The algorithm conservatively estimates the number of elastin layers by selecting the lower of the two median values, ensuring that over-counting does not occur. This approach accounts for any potential discontinuities or artifacts that might affect the layer count in one direction but not the other. Once the number of elastin layers is determined for each tissue region, it is stored in the elastin_layers attribute. The calculated number of elastin layers is appended to the elastin_layers list, which stores the number of layers for all tissue regions analyzed. If no elastin layers are found in a given region, the algorithm ensures that a default value of 0 is recorded. If no elastin layers are detected in any of the tissue regions, the algorithm ensures that the elastin_layers list is not left empty by setting it to [0]. This guarantees that even in the absence of elastin, the result is properly recorded as zero.

### Pipeline Outputs

The algorithm produces several outputs, both image-based (for visualization) and quantitative (for analysis and reporting). These outputs are generated during various stages of the analysis and can be saved for further examination or included in scientific reports. Image-based outputs are images generated at different stages of the analysis pipeline, providing a visual representation of the elastin structures and their characteristics. The algorithm generates four key types of images. The RGB image of the extracted region (based on the bounding boxes) is visualized and saved. This image represents the original tissue region before any processing or analysis. This serves as a baseline reference for understanding how the tissue looks in its raw state before any masks or networks are applied. The elastin thickness map is a grayscale image where each pixel along the elastin skeleton is assigned a value corresponding to the thickness of the elastin layer at that point. This map visualizes the variation in elastin thickness across the tissue and highlights areas where the elastin is thick or thin. It is useful for identifying potential structural abnormalities. The tortuosity map is a color-coded representation of the tortuosity values along the elastin network. Each pixel in the skeleton is assigned a tortuosity value, indicating how wavy or curved the elastin fiber is at that point. This map helps visualize areas where the elastin fibers are highly curved (higher tortuosity) versus relatively straight (lower tortuosity), which can reveal structural irregularities or pathologies in the elastic lamina. The elastin network map is a visualization of the network structure of the elastin fibers. In this map, pixels are color-coded based on their node type (tip points, branch points, or regular points). This map highlights the overall connectivity of the elastin network and shows how the fibers are structured. It allows for the identification of key features such as branch points and tip points. All the image outputs (RGB, thickness, tortuosity, and network) are saved to the specified output directory using the validation_viewer object. This allows for the creation of validation snapshots, which can be used to check the results visually and ensure that the processing steps were successful. Each image is saved with a specific filename that includes the prefix indicating the type of output (e.g., “elastin_thickness”) and the subsample number, ensuring proper organization and traceability of the results.

The algorithm also generates quantitative metrics, which summarize key aspects of the elastin structure, such as thickness, tortuosity, and the number of elastin layers. These metrics are calculated using various statistical methods and are stored in two key output structures: VVGStats and StatsHolder. The thickness stats summarize the distribution of elastin thickness across the tissue. These metrics are calculated from the elastin thickness map and include:

- Mean Thickness (thickness_mean): The average thickness of the elastin layers.
- Standard Deviation (SD) (thickness_sd): The variability in elastin thickness.
- Median Thickness (thickness_median): The central value in the distribution of thickness values.
- Quantiles (Q25, Q75): The 25th and 75th percentiles of the thickness values, giving an idea of the range of typical thicknesses.
- Minimum and Maximum Thickness (thickness_min, thickness_max): The extreme values in the thickness distribution.
- Coefficient of Variation (COV) (thickness_cov): The normalized variability of the thickness, calculated as the ratio of the standard deviation to the mean.
- Skewness (thickness_skew): A measure of asymmetry in the thickness distribution, indicating whether the distribution is skewed toward thinner or thicker values.

These statistics are stored in the StatsHolder object for thickness and are later used for reporting.

The tortuosity stats summarize the distribution of tortuosity values across the elastin network. These metrics provide insights into how wavy or curved the elastin fibers are. Key metrics include:

- Mean Tortuosity (tortuosity_mean): The average tortuosity across the entire network.
- Standard Deviation (SD) (tortuosity_sd): The variability in tortuosity values.
- Median Tortuosity (tortuosity_median): The central value of the tortuosity distribution.
- Quantiles (Q25, Q75): The 25th and 75th percentiles of the tortuosity values.
- Minimum and Maximum Tortuosity (tortuosity_min, tortuosity_max): The extreme tortuosity values across the network.
- Coefficient of Variation (COV): A normalized measure of the variability in tortuosity.
- Skewness: A measure of asymmetry in the tortuosity distribution.

These statistics are also stored in a StatsHolder object and used for reporting.

The number of elastin layers is a key output that describes how many distinct layers of elastin are present in each tissue region. This is calculated by analyzing the elastin mask within the tissue regions and counting the number of separate elastin components, as described in the previous section.

The network stats summarize key aspects of the elastin network, including the number of branches, tips, and the overall complexity of the network. Key metrics include:

- Number of Branch Points (num_branch_points): The total number of branch points in the network, where fibers diverge.
- Number of Tip Points (num_tip_points): The total number of tip points, where fibers end.
- Number of Trees (num_trees): The number of distinct connected components (trees) in the network.
- Segment-to-Network Ratio (segment_network_ratio): A ratio that reflects the continuity of the network. A lower value indicates more breaks, while a higher value suggests greater connectivity.
- Complexity (complexity): The overall complexity of the network, calculated as the sum of branch points, tip points, and trees.

All the key statistics for each subsample are collected and saved in a VVGStats object, which can be exported for further analysis or reporting. This includes the elastin thickness stats, tortuosity stats, network complexity, and the number of elastin layers. The VVGStats object contains all the key metrics for each tissue subsample, which can be written to a CSV file for easy analysis.

## Discussion

This study introduces a fully automated, stain-aware pipeline that converts Verhoeff–Van Gieson (VVG) whole-slide images of mouse aortae into reproducible, quantitative readouts of elastic-fiber architecture. By uniting optical-density deconvolution, robust skeletonization, and graph-based morphometry, the workflow delivers pixel-level maps of fiber thickness, lamina count, tortuosity, and network complexity—metrics that were previously obtainable only by labor-intensive manual tracing.

The ability to measure elastin thickness and continuity at scale sheds light on the mechanical reserve of the media, while local tortuosity highlights regions of incipient fiber fragmentation—an early event in aneurysm and hypertension models. Counting laminae provides an orthogonal index of medial organization that can be tracked across developmental stages, genetic backgrounds, or pharmacological interventions.

Collectively, these metrics enable powered genotype–phenotype screens in heterogeneous populations such as Diversity Outbred mice, and they furnish objective endpoints for therapies aimed at preserving or restoring matrix integrity.

Earlier methods typically (i) analysed cropped fields of view rather than whole slides, (ii) relied on color-threshold heuristics that fail when stains overlap, or (iii) treated elastic laminae as isolated objects, ignoring branching topology. Our pipeline overcomes each limitation. OD deconvolution disentangles elastin from collagen even in unevenly stained sections; multiresolution processing keeps gigapixel data tractable; and the graph representation preserves complex branching, allowing straightforward computation of continuity, segment-to-network ratio, and other network-level features.

Several factors could influence accuracy. First, segmentation thresholds—though empirically tuned—may bias thickness estimates in slides with extreme under- or over-staining. Second, tortuosity depends on the chosen search radius; excessively short radii underestimate waviness, whereas very long radii blur local curvature. Third, optical sectioning is two-dimensional; undulating laminae that dip in and out of the plane may be mis-classified as discontinuous. Future iterations could incorporate adaptive thresholding guided by deep-learning stain normalisation, multi-scale tortuosity measures, and 3D reconstruction from serial sections or micro-CT to mitigate these issues.

Machine-learning–based instance segmentation of elastin and smooth-muscle borders could unlock additional features such as inter-lamellar spacing or cell–matrix alignment^26–30^. Adapting the pipeline for three-dimensional image stacks obtained from serial sections^31^ or advanced imaging techniques like micro-CT could provide even more detailed structural information^30^. Extension to other elastin-rich tissues (lung, skin) or to different stains (e.g., Movat pentachrome) is straightforward given the modular design. Finally, integrating the pipeline with mechanical testing or µCT-derived strain maps would couple microstructural metrics to functional readouts, enabling true structure–function modelling of the arterial wall.

By providing quantifications of elastin area, thickness of lamina, the number of layers of elastin laminae, and tortuosity, this workflow facilitates investigations into the consequences of genetic variation on phenotypes related to the tissue structure and extracellular matrix composition of aortic tissue in mice. Genetic analysis of histology-based image-derived phenotypes in highly heterogeneous mouse cohorts, such as the Diversity Outbred (DO)^32^ or UM-HET3^33^ outbred populations, offers potential insights into the genetic underpinnings of aortic pathophysiologies including aneurysm and dissection^34^. Such analysis may prove complementary to genetic dissection of image-derived phenotypes in human cohorts in which tissue histology measurements are difficult to obtain. Furthermore, the methodology described herein can be applied to genetically modified animal models or experimental populations testing the effects of interventions on health outcomes, leading to mechanistic insights underlying diseases of the aorta.

By bringing whole-slide, quantitative rigor to elastin histopathology, the proposed framework bridges a persistent gap between qualitative microscopy and biomechanical phenotyping. Its scalability, openness, and rich output suite make it a practical foundation for large-scale vascular studies and a potential supporting tool for future diagnostic workflows.

## Methods

### Programming Environment

The ROI extraction process and analysis pipeline was implemented in Python using libraries such as NumPy for numerical operations, SciPy for image filtering and morphological operations, scikit-image for image processing functions, and tifffile for reading multi-resolution TIFF images.

## Code availability

The pipeline is fully written in Python. The Python code is open-source and freely available online via Github at https://github.com/aelefebv/histo-aorta-features.

## Contributions

A.E.Y.T.L. conceived of and designed the pipeline, wrote the manuscript, created the figures, developed and wrote the code, performed the data analysis, and supervised the study. M.M. provided background and motivation for the creation and refinement of the pipeline. M.M. provided helpful feedback on experimental design, data interpretation, and edited the manuscript.

## Acknowledgments

We thank Graham Ruby for helpful discussions on the pipeline and its applications, and for providing much of the motivation for this paper, as well as the samples used for the creation and testing of this pipeline. We also thank Calico Life Sciences for supporting this work.

